# FAIR sharing of molecular visualization experiences: from pictures in the cloud to collaborative virtual reality exploration in immersive 3D environments

**DOI:** 10.1101/2020.08.27.270140

**Authors:** Xavier Martinez, Marc Baaden

## Abstract

Motivated by the current Covid-19 pandemic that has spurred a substantial flow of structural data we describe how molecular visualization experiences can be used to make these datasets accessible to a broad audience. Using a variety of technology vectors related to the cloud, 3D- and virtual reality gear, we examine how to share curated visualizations of structural biology, modeling and/or bioinformatics datasets for interactive and collaborative exploration. We discuss F.A.I.R. as overarching principle for sharing such visualizations. We provide four initial example scenes related to recent Covid-19 structural data together with a ready-to-use (and share) implementation in the UnityMol software.

**Synopsis:** Visualization renders structural molecular data accessible to a broad audience. We describe an approach to share molecular visualization experiences based on FAIR principles. Our workflow is exemplified with recent Covid-19 related data.

## 1. Introduction

The Covid-19 pandemic has spurred a wealth of new structural biology data related to the severe acute respiratory syndrome coronavirus (SARS-CoV)-2 retrovirus and its interactions with other macromolecules involved in the development and spread of the disease [1, 2]. Such structural data is transparent to experts in the field, but requires adequate visualization to become accessible to a broader audience. As put by Card et al., “visual artifacts aid thought; in fact, they are completely entwined with cognition action” [3]. Visualizations prepared by experts may serve both educational and research purposes by highlighting key aspects of a given dataset or comparing features among several ones thereby supporting thought and reasoning. Here we aim to establish a general workflow to prepare such curated educational and research material, taking the advent of recent Covid-19 structural and modeling data as example. Our overall goal is to enable easy sharing of a given visual experience with others so that the same content can be accessed in various ways. We put focus on collaborative virtual reality (VR) as well as sharing 3D models and scenes to intuitively convey the spatial complexity of these molecular objects. Shape-related features bear particular relevance for drug design and hence for expert users, yet they can also be understood by an inexperienced person, in particular through a visual experience such as those described here. A particular aim is to treat the sharing of visual experiences with the same F.A.I.R. principles [4] as one should apply for the underlying raw chemical [5], structural and modelling data, and as is generally the case in crystallography [6]. This target is challenging by itself, as common FAIR-based sharing platforms do not provide a category that is well adapted to visual experiences.

The features of the workflow we are seeking to implement are schematically summarized in Figure 1. Starting from the original raw data that should encompass experimental structural datasets, molecular modelling results and bioinformatics findings, we aspire to prepare new shareable visual content to be set up and curated by an expert. To do so, we will use a molecular visualization software package to describe the visualization scene and all its annotations through command scripts. The experience can be customized through add-ons, for example providing dataset-specific user menus leading to a visual scene that can be explored directly in the software. This exploration does however require learning at least some basic usage of that software and how to manipulate the scene. This task can be simplified to some extent by designing appropriate simple-to-use custom user menus. As an alternative, the core content to be visualized can be exported to a variety of media such as still images, movies of various formats – possibly stereoscopic ones –, native 3D models or a fully interactive scene to be explored with advanced technology. We aim to support a broad variety of sophisticated hardware such as VR headsets, wall-size stereoscopic displays and the HoloLens, but also simple setups such as Google cardboard or even just running in a web browser. To render these media findable, accessible, interoperable and reusable, a variety of options are discussed.

**Figure 1.**
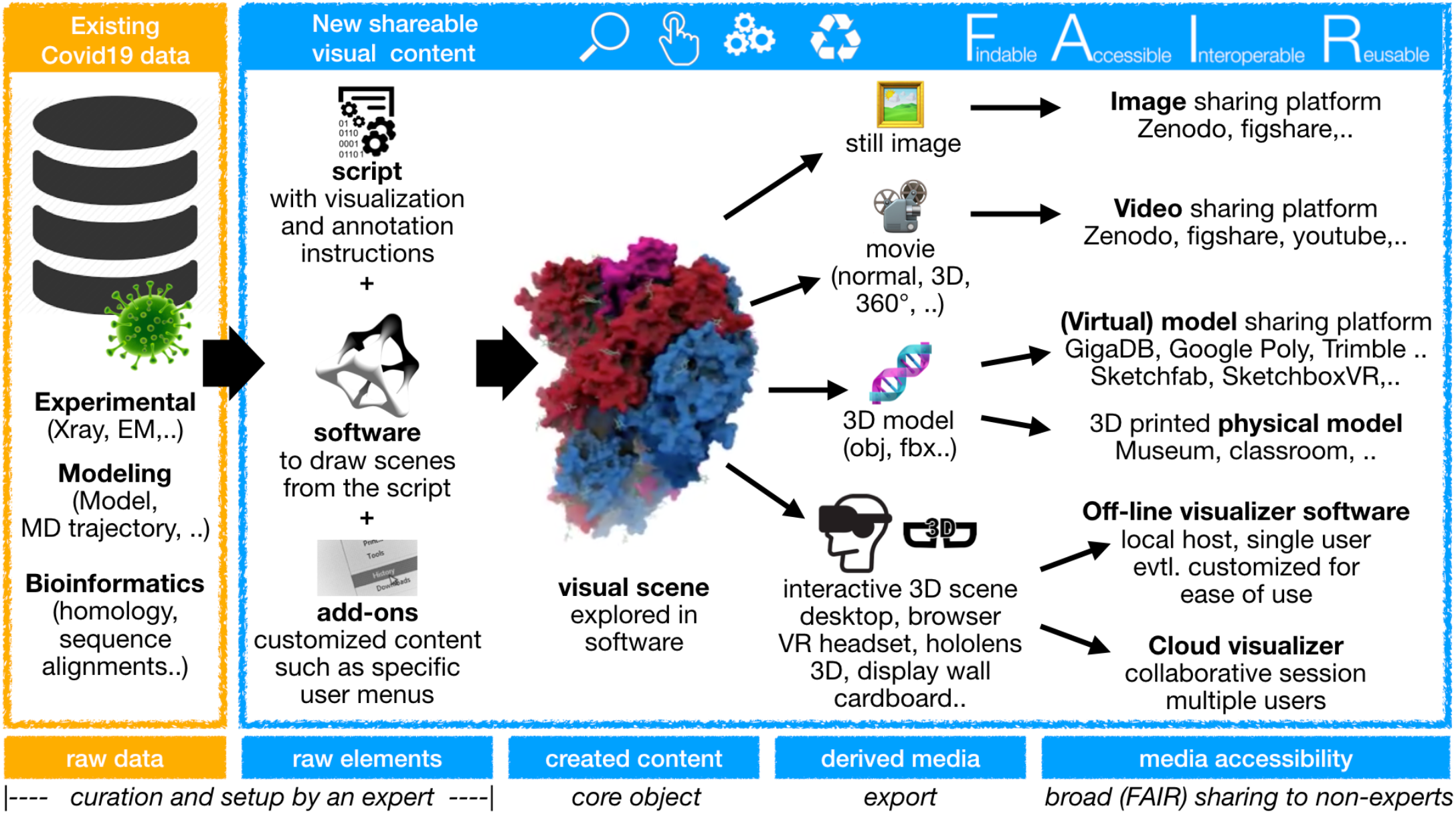
A scheme summarizing the suggested workflow for FAIR sharing of molecular visualization experiences, building on Covid-19-related structural data. Using existing data, new shareable visual content is created from raw elements such as a script with instructions, software to execute the script and eventual add-ons to facilitate the end-user tasks (for instance custom menus to provide only the strictly necessary shortcuts for a given experience). The visual scene that is created can be explored within the software itself or it can be exported in various ways to generate derived media such as images, movies, 3D models and even full experiences in e.g. VR contexts. These media can be rendered accessible through platforms that implement FAIR principles and ease their exploration.

The set of (open source) software plus openly accessible scripts that we propose to provide make the visual experience as reproducible as possible [7]. Inherent limitations arise from technical hardware and software dependencies related to rapidly evolving visualization technology that are by nature short-lived with a high rate of change. We chose the UnityMol package to implement our proof of concept set of visualization experiences, as we have been developing and redesigning this software, which provides specific advantages for the tasks at hand in its latest version [8, 9]. Many alternative software options of high value could have been considered such as those recently reviewed for protein visualization [10]. We opted for UnityMol as it provides a good compromise between feature-richness and ease of use. A wide variety of media and technology vectors is supported, while the software remains simple to use for a broad range of users due to its single-window design with only a few menus to operate.

Our ambition is to provide a few proof of concept implementations that others can build on subsequently to further improve or simply adapt them. We have implemented all functionalities that we deemed necessary to prepare the Covid-19-related materials, and present here four basic examples. Based on these example scenes, we experiment with applying F.A.I.R. principles to such visualization-centric data. One way to achieve this goal is by addressing the setup of the visualization experiences, i.e. the underlying software and the related scripts (designated as “raw elements” in Fig. 1), rather than the resulting media products. A more classical approach is to provide the derived media, which may consist in videos or images for which sharing platforms exist, although this is not enough to optimize FAIR compliance [11].

Sharing of visualization experiences appears as a particularly timely topic. The case of the Covid-19 pandemic provides a stringent example for the need to render the latest research data explorable by the broad scientific community. If such data were only to be provided in its raw form, the accessibility would be quite limited. The worldwide Protein Data Bank [12] indeed provides quite a few visualizations along with the actual data files. Recently, the european PDB also proposed a visual analytics platform that is FAIR compliant [13]. Here we try to go one step further in the democratization of such data by preparing curated visual experiences as a valuable support for cross-disciplinary scientific exchange. In related fields, similar initiatives have recently emerged. Now more than ever, sharing data from molecular simulations and modeling is becoming a priority to ensure reproducibility and accelerate discovery [14]. An increasing number of initiatives aim to develop open access and reproducible molecular simulations (see for example [15–17] among other approaches).

Sharing such molecular dynamics (MD) simulation data equally needs efficient visualization and collaboration tools [18] for apprehending such data remotely to avoid unnecessary data transfers. Yet to the best of our knowledge, current initiatives focus on the actual data, not so much on the visualizations and annotation thereof. Concerning the specific case of Covid-19, many data portals and hubs were put together to provide more unified access to relevant data. A brief selection is listed in Table 1 and served as testing ground for our visualization experiments. Many of the listed portals provide a very direct access to the data we require for our visualization experiences, a few provide more general search engines with a less direct path to the actual dataset of interest. We did not include the classical routes to search data such as the PDB, GigaDB [19, 20] or Zenodo and we only included one of the many relevant initiatives by individual scientists or laboratories as example.

**Table 1.**
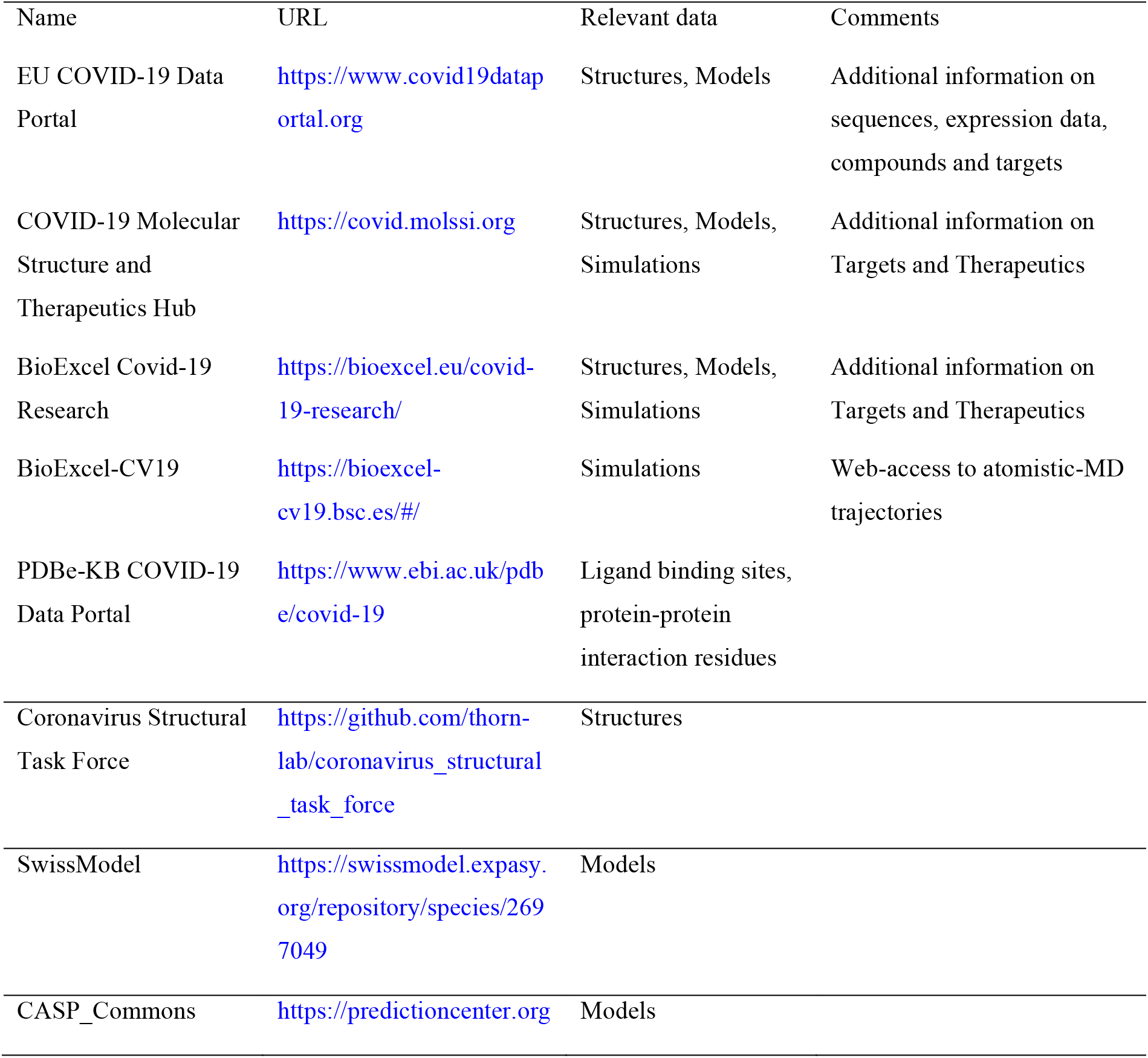
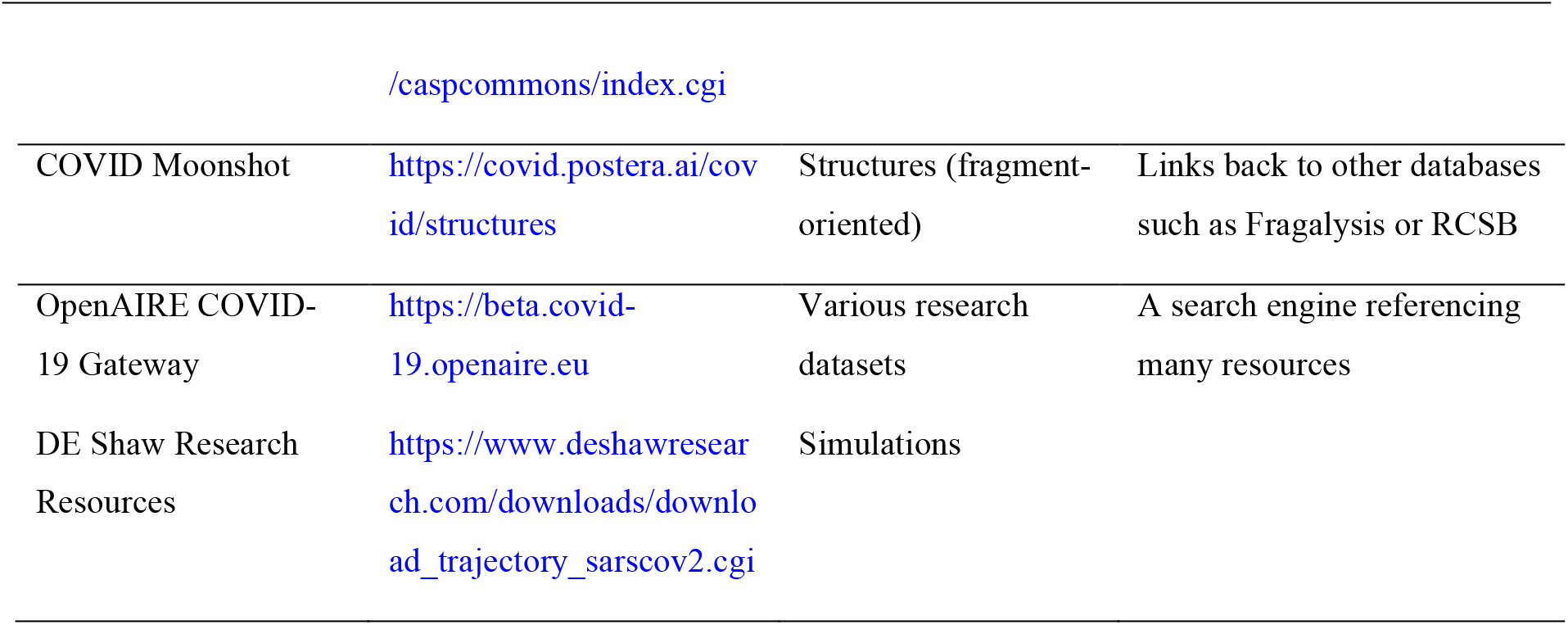
A few examples of Covid-19 data portals. Some of the data in these portals are likely to be cross-referenced or duplicated. The selection of portals is by no means exhaustive, with the number of available resources rapidly increasing.

## 2. Materials and Method

Here we will briefly describe key functionalities implemented for the purpose of sharing molecular visualization experiences. As an overarching principle, we want to be able to generate a broad variety of media, including still images and videos based on these visual scenes describing the core of the contents, capturing them from within the running software, ideally in an executable build and for a handful of technically more involved features from within the Unity3D editor environment. For this purpose we implemented, extended and stabilized several features. A certain number of those features are still experimental only.

We added a python console and API for scripting within UnityMol, providing an easy way to formalize and share the narrative of a given experience, as well as facilitating the reproducibility of all necessary steps leading to a scene. We furthermore implemented the possibility to generate custom user menus so that complex and fine-tuned functionality can be achieved with the click of a button. A possible practical use could be to group shortcuts to relevant structural ensembles in light of Covid-19 for visual analysis, comparison or even manipulation. The latter can be achieved through interactive simulations: UnityMol implements the generic IMD protocol for steering molecular simulations through MDDriver [21], and provides a few in-app implementations such as a rigid protein-protein docking module.

The scenes can be annotated by the user through several functionalities such as text bullets or free hand drawing. A guided tour function was implemented to walk the user through a set of pre-determined key frames in a fully interactive experience.

A particular focus concerns the possibility to generate effectful high-quality graphics, for instance in order to convey the complex shapes of molecular surfaces using ambient occlusion (AO) or to draw the user’s attention to a certain part of the structure of interest using photography effects such as depth-of-field (DOF). In addition to the molecular structures themselves, their properties such as electrostatic field can be depicted as well [9]. In the latest version of the software we added an experimental feature enabling interactive raytracing using the OSPRay [22] package. This is illustrated for our first example of the spike trimer complex with the ACE2 enzyme (Figure 2A). We added a custom menu to simplify navigating through this example (Figure 2B). We explored the scene collaboratively in a 3-user virtual reality session (Figure 2C) where avatars represent each participant. To capture such sessions, still images from screenshots or movies from video capture can be created at user defined resolution. These media can be generated and exported with specific options such as a 360 degree view or a 3D stereoscopy enhancement. For these purposes, specific cameras had to be added within Unity. In general, image and video exports do however not preserve the full complexity of the initial object. For this purpose we added the export of textured 3D polygonal objects in the common obj, fbx and probably soon also glTF formats. Special effects such as AO and DOF cannot currently be exported. Such 3D models can be transformed into tangible physical objects through 3D printing [23].

**Figure 2.**
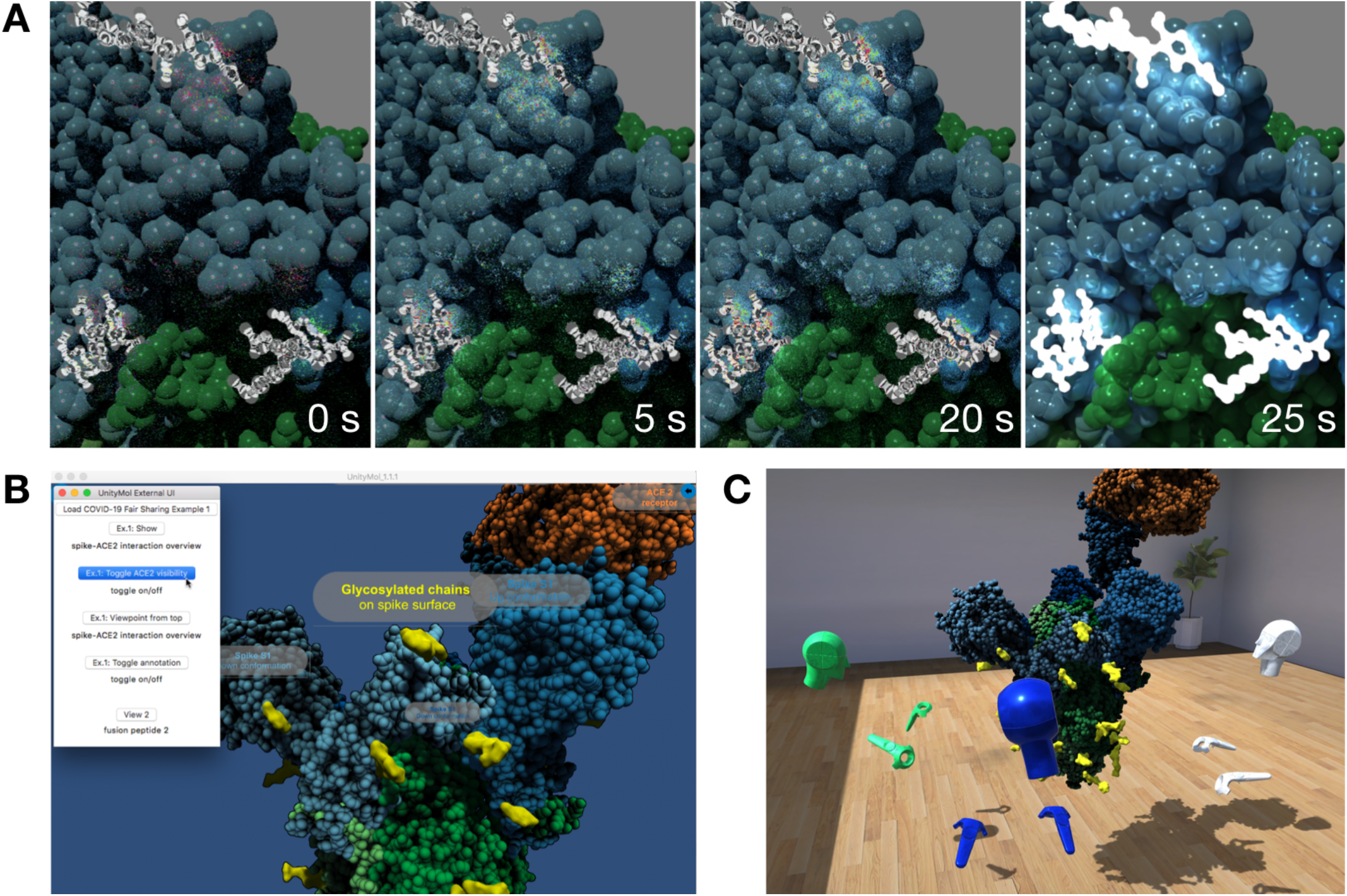
Ilustration of three types of visual experiences within UnityMol. **A)** Interactive raytracing of a simple structural view of the spike-ACE2 complex (example 1). A series of four images at distinct time intervals is shown to depict the ongoing work of the raytracer. After 25 s a denoiser pass is executed, significantly increasing the visual clarity of the scene. The raytracer continues indefinitely to improve the scene. The fluidity and interactivity are only limited by the CPU-bound performance. Indicated timing was obtained with a 2013 3,5 GHz 6-Core Intel Xeon E5. **B)** A custom menu extension for example 1, providing the possibility to toggle annotations on/off along with direct access to different representations and viewpoints. **C)** Screenshot of a collaborative multi-user session where 3 participants are examining example 1 with virtual reality headsets and controlers.

## 3. Results

We have started the process of building up a collection of Covid-19 example scripts with several objectives: they should be easy to understand, even for non-specialists; others should be able to re-run them under various conditions in terms of visualization hardware and modality and (in the future) it should be simple to use them as starting scene for shared multi-user sessions. A particular emphasis lies on the design of visual experiences suitable and tested for virtual reality (VR) exploration as our current research is focused on this technology. In our opinion its potential is still largely under-explored in academia and industry, be it for research, collaboration or education purposes [24]. To illustrate the sharing of such visual experiences, we have prepared four typical datasets depicted in Figure 2 and Figure 3): a simple structural view, a collection of small molecules binding to a viral protein, an MD trajectory and a bioinformatics sample with conservation as well as mutation data.

**Figure 3.**
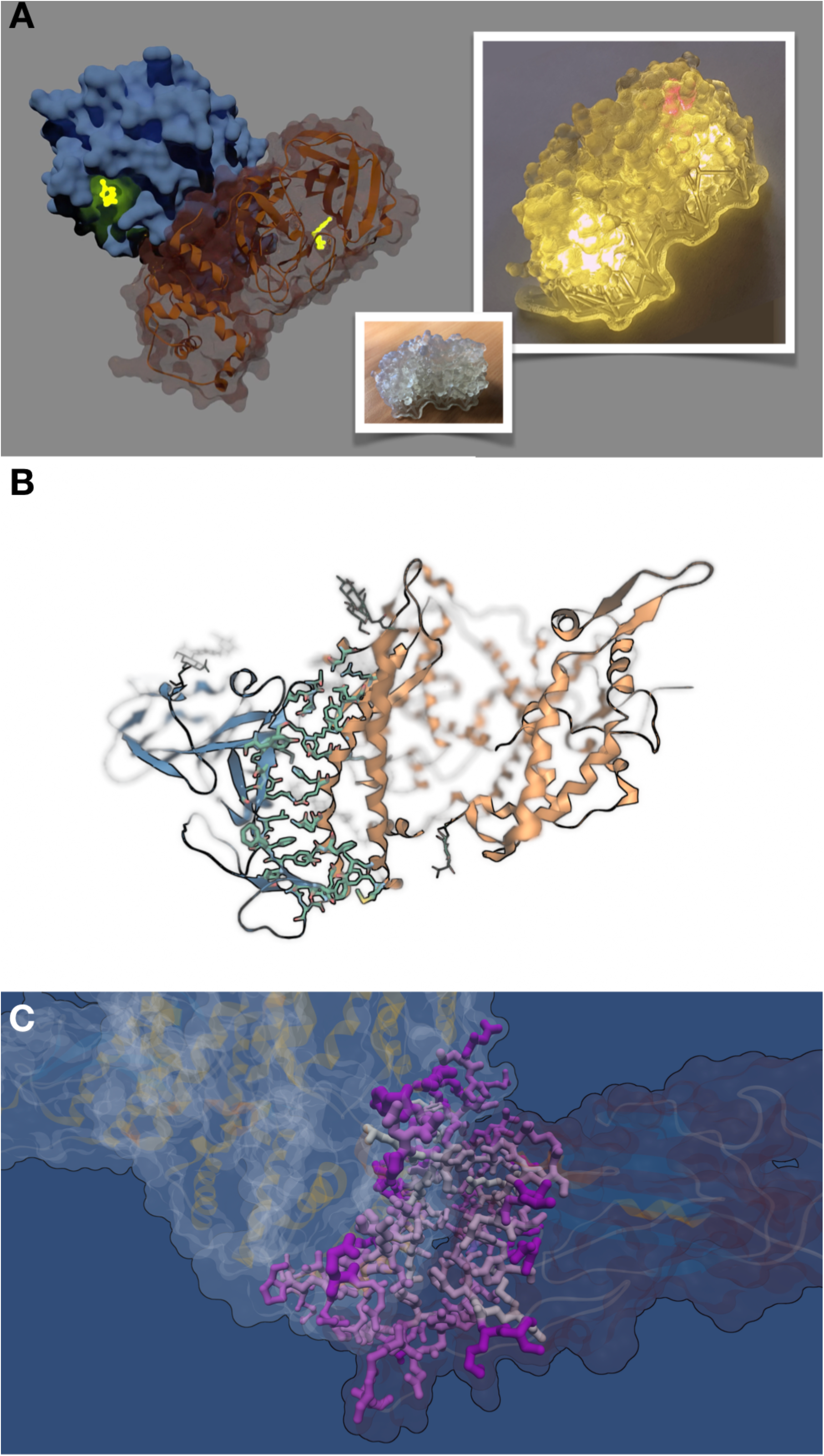
A glimpse into examples 2 to 4. **A)** Example 2 depicts drug molecules (shown in glowing yellow) binding to the dimeric biological assembly of Covid-19 main protease (monomers shown in blue surface and brown cartoon) in 92 protein databank structures. The image is a snapshot taken from an animation of the whole dataset. The insets show a 3D printed translucent monomer model (bottom) and how one may illuminate it (right) to highlight specific areas with a laser (red spot). The orientation of the right image is similar to the one of the brown monomer subunit shown on the left. **B)** Example 3 renders a molecular dynamics simulation of the SARS-CoV-2 spike receptor-binding domain binding to the ACE2 receptor [25]. A depth-of-field effect is used to draw attention to the binding interface. **C)** Example 4 represents results on species variability of the binding interface from a bioinformatics analysis. The interacting residues are colored in a pink color scale according to the Haddock docking score from [26] and their size is varied according to residue conservation assessed through a Shannon entropy measure.

The first example depicted in Figure 2 aims at setting up a simple structural view of the SARS spike glycoprotein complex with human angiotensin-converting enzyme 2 (ACE2). The trimeric complex accessible from PDB-id 6CS2 [27] was used as starting point for many early SARS-CoV-2 studies.

We then have a comparative look at where drug molecules bind to the main protease of Covid-19 in our second example (Figure 3A). The visualization is inspired by the animation of small molecules in 92 protein databank structures (https://www.rbvi.ucsf.edu/chimerax/data/sars-protease-may2020/) prepared by the ChimeraX [28] team. We created a 3D print of the protease monomer as well.

Molecular dynamics simulations provide another inestimable resource for insight into molecular mechanisms. Example 3 (Figure 3B) is based on a trajectory depicting a binding event of the receptor-binding domain (RBD) of the SARS-CoV-2 spike and the human ACE2 receptor (DESRES-ANTON- [10857295,10895671] of ref. [25]).

The fourth example illustrates a visual experience related to representing the results of bioinformatics analyses. Our example is built upon the freely available data from a recent study on cross-species transmission of SARS-CoV-2, highlighting the species variability of viral-host protein interactions [26]. We visually map this data onto the ACE2-RBD complex (Figure 3C).

The first level of sharing these four examples is to make the software and the scripts themselves available. We considered the zenodo and GigaDB platforms and chose zenodo as main platform for our experiment. Typically, no dedicated category for “visual experiences” exists in these databases, whereas the classical ones such as software, dataset, image, video, workflow or other only partially reflect this case. As a first approach, we created a Zenodo community (Figure 4A) on FAIR sharing of molecular visualization experiences at https://zenodo.org/communities/fair-molvisexp/. We then linked a github repository for our visualization scripts to Zenodo and the collection. We produced a set of derived materials including pictures, movies and 3D objects. Cloud-platforms for pictures and movies are very common nowadays and we will not go into much detail on this aspect. We used both the zenodo and figshare platforms (http://figshare.com) for pictures and for movies, the former one for consistency and the latter because of its visually oriented interface and popularity. To regroup our productions on figshare, we created a collection (https://doi.org/10.6084/m9.figshare.c.5101400) shown in Figure 4B. We also experimented with tangible physical objects as a particularly accessible form for 3D models through 3D printing. Many generic 3D printing databases exist, whereas the NIH 3D print exchange [29] is a specific research initiative dedicated to bioscientific 3D prints. We deposited our models in this resource (Figure 4C). Such 3D models can also be disseminated and shared virtually, which is a less common process. We experimented with four platforms: Sketchfab (https://sketchfab.com; Figure 4D), Google Poly (https://poly.google.com; Figure 4E), SketchBoxVR(http://sketchboxvr.com)andTrimble3Dwarehouse (https://3dwarehouse.sketchup.com). For the time being we used the dataset, software, figure and video categories to reference all information on Zenodo. We have created a versioned catalogue page ofalltheoutcomesoftheseexperimentsat https://github.com/bam93/fair_covid_molvisexp/tree/master/overview. Once all the functionalities described here are stabilized in the UnityMol software, we will provide instructions on how to export to these platforms on the same catalogue page.

**Figure 4.**
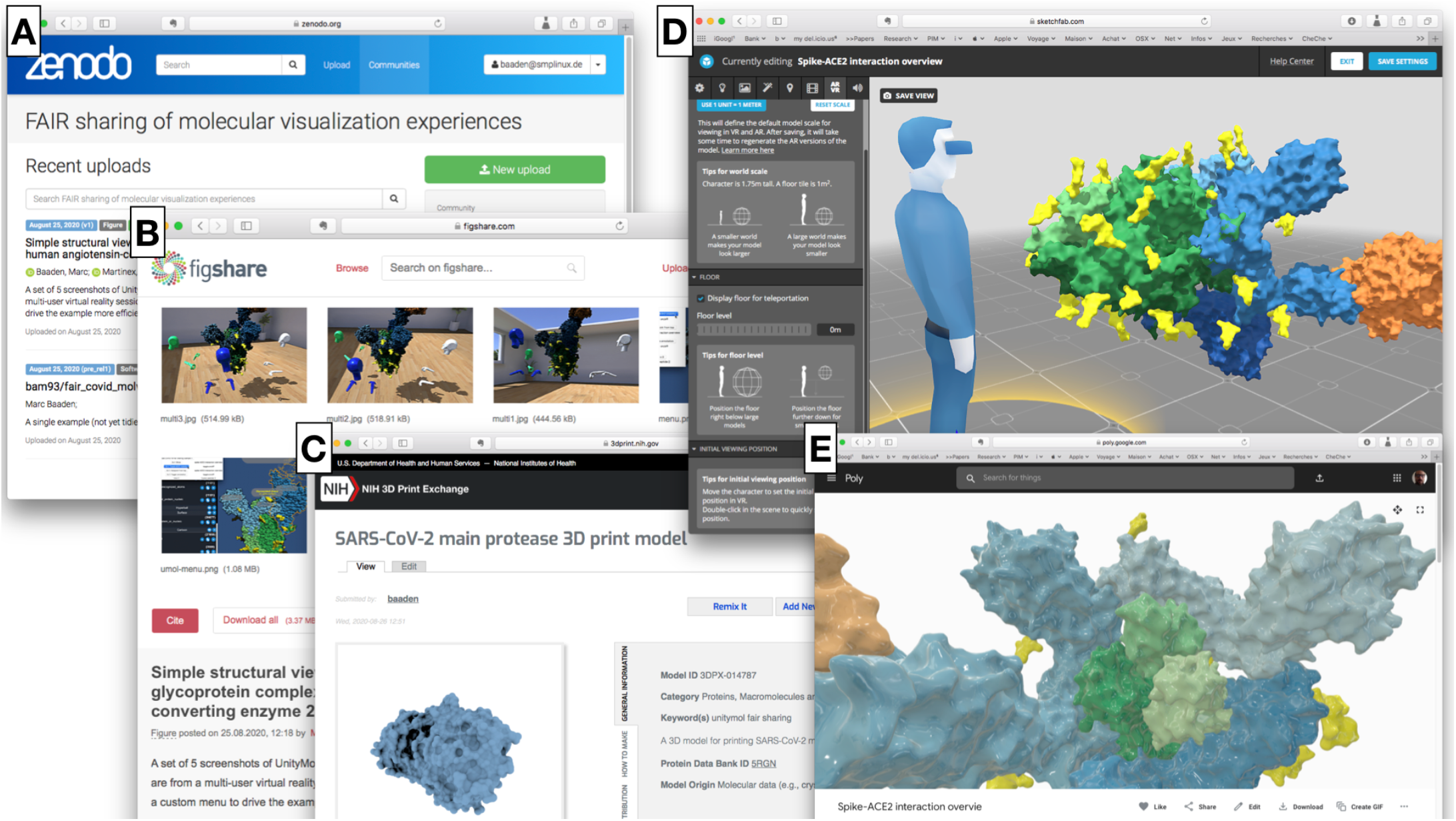
A visual overview of content sharing platforms hosting our visual experiences to make them FAIR-compliant. **A)** depicts a zenodo community regrouping all the content created for this work (https://zenodo.org/communities/fair-molvisexp/). **B)** is a figshare collection of a subset of this content (https://doi.org/10.6084/m9.figshare.c.5101400). **C)** illustrates the entry of our 3D printed model from Fig. 3A on the NIH 3D print exchange (https://3dprint.nih.gov/discover/3DPX-014787). **D)** illustrates a 3D model uploaded to Sketchfab (https://skfb.ly/6UFOw) with the possibility to edit AR and VR viewing settings, as the platform automatically enables access to mobile phone and desktop virtual reality rendering in addition to web-based exploration. **E)** shows the same 3D model rendered in the Google Poly web-based explorer (https://poly.google.com/view/5zsJiglTWbm). **C, D** and **E** are accessible through the respective referenced 3D model datasets on zenodo, an stl file for 3D printing and an fbx file for 3D model building.

Referencing our datasets on these platforms only represent the first, relatively easy step towards implementing the FAIR principles. The main issues may lie in the interoperability and reusability aspects for the (meta)data associated with the visual experiences. For example, metadata is needed to describe software version, script purpose, underlying datasets, dependencies and the variants of output that can be produced. There are definitely many ways to go about implementing the FAIR principles. In this early attempt at visual experiences, we first and foremost want to raise awareness about this yet overlooked category, but do not ambition to provide a full-fledged solution. Concerning the fact that our data should be Findable, the choice to target Covid-19 makes this goal relatively easy to achieve as there is an abundance of hubs for increasing the findability of such data as described briefly above (Table 1). The Accessible attribute should be taken care of by our choice of well-established FAIR-compliant databases for most of the produced media. Interoperability is clearly an area for further improvement. In particular we consider bridges between different visualization environments as an important way forward. Initiatives fostering universal protocols that can be interchanged such as MolQL for the query language to define selections [30] need to be extended to describe all ingredients needed for a visual experience, in particular the molecular scene representation. As a modest step in this direction we have experimented with a Pymol session interpreter for Unitymol (https://github.com/LBT-CNRS/PymolToUnityMol), so that Pymol users can re-use their previously prepared scenes. Reproducibility is intrinsically a difficult aim for visual experiences. However, by providing scripts with full instructions for the setup of the experience and by making the corresponding visualization software available as versioned open source project along with ready-to use executable builds for many operating systems and platforms, reproducibility and accessibility are maximized. Furthermore, the software is conceived within a game engine, which by design is easily extensible, with a shallow learning curve.

## 4. Outlook

From this early experiment, we plan to expand the range of example visual experiences, be it based on the context of the rapidly growing Covid-19 data or more generally the overwhelming structural, modeling and dynamic data of macromolecular assemblies. We are trying to optimize the generated objects so that they can run on devices with limited hardware specifications. An example would be the HoloLens, with good ergonomy but limited graphical power [31] for which we want to be able to export simplified FBX/gltf polygonal models to run smoothly. Most of our efforts will be dedicated to refine the multi-user collaborative features to enable visual sessions for joint exploration.

More generally, our contribution calls for considering whether visual experiences should be assigned a specific category to be added to FAIR sharing platforms in order to take into account the specificities of such interactive, computer-supported graphical representations of research data.

## Acknowledgements

This work was supported by the “Initiative d’Excellence” program from the French State (Grant “DYNAMO”, ANR-11-LABX-0011 and grant “CACSICE”, ANR-11-EQPX-0008). X.M. and M.B. thank Sesame Ile-de-France for co-funding the display wall used for data analysis. X.M. and M.B. thank UCB Biopharma for support. We thank Nawel Khenak for help with 3D printing and Nicolas Férey and Antoine Taly for experimentations with laser and LED-lighting of translucent 3D-printed molecules.

